# Pseudotime analysis of 2,106 brains across nine regions reveals conserved immune, neuronal, and myelin regulatory programs in Alzheimer’s disease

**DOI:** 10.64898/2026.08.18.745353

**Authors:** F Ecca, S Song, M Naymik, MJ Huentelman, IS Piras

**Affiliations:** Early Detection and Prevention Division, Translational Genomics Research Institute

**Keywords:** Sporadic Alzheimer’s Disease, *post-mortem* tissue, multiple brain regions, RNA expression, pseudo-time trajectory (PT)

## Abstract

Pseudotime trajectories can reconstruct latent disease progression from cross-sectional transcriptomic data. However, whether Alzheimer’s disease (AD) progression follows a conserved molecular architecture across brain regions remains unclear. We applied pseudotime analysis to harmonized bulk RNA-seq data from 2,106 postmortem brain samples (1,364 AD, 742 controls) across nine brain regions from three AMP-AD cohorts (ROSMAP, Mayo, MSBB). Pseudotime was significantly associated with AD diagnosis in all nine regions and with Braak stage in seven of nine. We identified 21 genes with concordant pseudotime associations across all regions, increasing to 234 when the cerebellum was excluded. Pathway analysis revealed 1,268 significant associations, with synaptic deregulation as the most conserved process, and immune/ECM programs showing greater regional specificity. The cerebellum followed a distinct pattern, with enrichment for protein refolding and chaperone pathways rather than neurodegeneration. Co-expression network analysis identified six conserved metamodules, including immune/glial (MM1) and excitatory neuronal (MM2) programs spanning all nine regions, and an oligodendrocyte/myelin program (MM3) in seven cortical regions. Key driver analysis identified 76 unique genes across 35 modules, with *HCK* and *LAPTM5* as the most broadly replicated immune regulators in seven regions. Oligodendrocyte-associated key drivers (*MYRF*, *CNP*, *MOBP*) increased along pseudotime in cortical regions, supporting active myelin remodeling during AD progression. These findings reveal a conserved transcriptional architecture underlying AD progression, organized around coordinated immune activation, synaptic loss, and myelin remodeling, with the cerebellum following a distinct trajectory.

## Introduction

Sporadic Alzheimer’s disease (sAD), represents a growing global health and socioeconomical burden^1^, as the number of people living with dementia is projected to increase from 57 million cases in 2019 to 153 million by 2050^2^ and the macroeconomic burden of Alzheimer’s disease and other dementias is estimated at $14.5 trillion between 2020 and 2050^3^. Its heterogeneity, which spans age at onset, sex, genetic risk, and environmental interactions, poses a major challenge for understanding disease mechanisms and developing effective therapies. While transcriptomic analyses have revealed key molecular pathways, cellular alterations, and gene networks associated with distinct genetic backgrounds, including APOE variants in sporadic disease and *PSEN1*-E280A mutations in autosomal dominant AD, a dynamic, unified view of disease progression remains elusive^4–6^. Most transcriptomic studies are cross-sectional, capturing static snapshots that fail to resolve the continuous evolution of molecular and cellular changes. As a result, critical aspects of disease trajectory and temporal ordering of pathological events remain poorly understood. Trajectory inference methods, such as pseudotime analysis^7–10^, offer a solution by reconstructing latent progression directly from gene expression profiles, providing a data-driven approximation of disease staging independent of clinical labels. This is particularly valuable in sAD, where clinical diagnosis is coarse and neuropathological staging provides only a single, terminal observation per individual. Although widely used in single-cell studies^11–13^, trajectory inference has been comparatively less explored in bulk RNA sequencing data from large, heterogeneous human cohorts.

Previous studies have demonstrated the feasibility of reconstructing molecular trajectories from bulk transcriptomic profiles of post-mortem AD brains. Mukherjee et al. used manifold learning to derive a molecular disease pseudotime from bulk RNA-seq data from the ROS/MAP and Mayo cohorts, identifying strong associations with Braak stage, CERAD score, and cognitive diagnosis^14^. A subsequent deep learning-based study derived a pseudo-temporal trajectory from brain transcriptomic data and showed that the resulting severity index captured neuropathological and clinical measures of AD progression across independent cohorts and brain regions^15^. However, whether transcriptional trajectories of AD progression exhibit a conserved molecular architecture across multiple anatomically distinct brain regions remains unclear.

AD pathology is also region-specific, with distinct brain areas showing differential vulnerability and temporal involvement^15,16^. Capturing both shared and region-specific transcriptional trajectories is therefore crucial for a comprehensive understanding of disease progression and for identifying conserved versus regionally restricted molecular programs. Therefore, if a common latent axis of disease progression exists, it should be detectable independently in multiple regions and should order samples consistently with neuropathological burden.

In this study, we applied a Bayesian latent variable model fit by variational inference^7^ to harmonized RNA sequencing data from the AD Knowledge Portal, encompassing nine brain regions from human postmortem samples collected in the Religious Orders Study/Memory and Aging Project (ROS/MAP), Mayo Clinic, and Mount Sinai VA Medical Center Brain Bank (MSBB)^17–19^. Specifically, we analyzed dorsolateral prefrontal cortex (DLPFC), anterior cingulate cortex (ACC), posterior cingulate cortex (PCC), temporal cortex (TCX), cerebellum (CBE), frontal pole (FP), superior temporal gyrus (STG), parahippocampal gyrus (PHG), and inferior frontal gyrus (IFG) for a total of 2,106 post-mortem brains (AD = 1,364, ND = 742). By reconstructing pseudotemporal trajectories, we aim to reveal transcriptional unique e shared programs associated with disease progression across anatomically distinct regions.

## Methods

### Datasets and samples description

We analyzed harmonized RNA-Seq data from postmortem brain samples obtained from the Religious Orders Study/Memory and Aging Project (ROSMAP), Mayo Clinic Brain Bank (MAYO) and Mount Sinai Brain Bank (MSBB) cohorts (accession number: syn21241740). The ROSMAP data include samples from the dorsolateral prefrontal cortex (DLPFC), anterior cingulate cortex (ACC), and posterior cingulate cortex (PCC). The Mayo Clinic cohort data consist of samples from the temporal cortex (TCX) and cerebellum (CBE), while the MSBB cohort data are sampled from the frontal pole (FP), superior temporal gyrus (STG), parahippocampal gyrus (PHG), and inferior frontal gyrus (IFG). Details regarding clinical data and the handling and processing of postmortem samples in ROSMAP^17^, Mayo^18^, and MSBB^19^ have been previously published^20^. Sample size and phenotype information are reported in **Table 1**. Diagnosis was harmonized across studies by standardizing the definition of Late-Onset AD (LOAD) to require both clinical and neuropathological confirmation. LOAD cases exhibit high neurofibrillary tangle and amyloid plaque burdens along with cognitive impairment, while controls (CTL) show low levels of pathology and preserved cognitive function (**Table S1).**

**Table 1:** Sample size and characteristics of the dataset used in this study. Data were downloaded after approved request from Synapse.org and are part of the RNAseq Harmonization Study (accession number: syn21241740).

| Study | Brain Region | AD | ND | Tot |
| --- | --- | --- | --- | --- |
| MAYO | Cerebellum (CBE) | 79 | 65 | 144 |
| MAYO | Temporal Cortex (TCX) | 80 | 68 | 148 |
| MOUNT SINAI | Frontal Pole (FP) | 146 | 69 | 215 |
| MOUNT SINAI | Inferior Frontal Gyrus (IFG) | 132 | 69 | 201 |
| MOUNT SINAI | Parahippocampal Gyrus (PHG) | 155 | 71 | 226 |
| MOUNT SINAI | Superior Temporal Gyrus (STG) | 156 | 64 | 220 |
| ROSMAP | Anterior Cingulate Cortex (ACC) | 171 | 94 | 265 |
| ROSMAP | Dorsolateral Prefrontal Cortex (DLPFC) | 294 | 143 | 437 |
| ROSMAP | Posterior Cingulate Cortex (PCC) | 151 | 99 | 250 |

### Preprocessing and pseudotime trajectories

Normalized and residualized gene expression count matrices were downloaded from the RNAseq Harmonization Study of the AD knowledge portal (syn21241740). Upstream processing was performed by the AMP-AD consortium using the RNAseq Reprocessing Workflow. In brief, BAM files were sorted with *Picard v2.2.4 SortSam* and converted to FASTQ with *Picard SamToFastq*; reads were realigned to the GRCh38 reference genome (GENCODE release 24) with *STAR v2.5.1b* using *twopassMode basic*, and gene-level counts were generated with the *STAR --quantMode GeneCounts* option. Genes expressed above 1 CPM in at least 50% of samples within each tissue and diagnosis category, and for which gene length and GC content were available from the *BioMart* December 2016 archive, were retained. Raw counts were normalized and adjusted for covariates by the consortium following the framework of Sieberts et al.^21^. Briefly, conditional quantile normalization was applied to account for gene length and GC content, followed by weighted linear modelling with the *voom–limma* pipeline; outlier samples were removed where PCA and hierarchical clustering agreed; known covariates were selected per study by stepwise weighted fixed/mixed-effect regression, and hidden confounders were estimated by iteratively re-weighted surrogate variable analysis. Expression was then modelled as a function of diagnosis, sex, study-specific covariates and surrogate variables, with donor as a random effect and observation weights derived from the *voom–limma* pipeline, fit separately within each study. Covariates and surrogate variables were selected independently for each study and brain region. For our analyses, we used the residualized matrices in which surrogate variables, sex and age at death were regressed out and only the diagnosis coefficient was added back.

We annotated gene symbols using *BioMart*, and we excluded genes lacking annotation for HGNC symbol, Entrez ID or Ensembl ID. We estimated the median absolute deviation (MAD) retaining only genes in the top 50% of the distribution. Pseudotime trajectories were independently estimated by study for each brain region using the *phenoPath* method^7^. The following settings were used: *elbo_tol* = 1 × 10^-05^, *z_init = “random”*, *thin = 10*, and *maxiter = 10,000*. ELBO plots were inspected to assess convergence. The relationships between pseudotime trajectories and clinical or neuropathological variables were evaluated using Pearson’s correlation, while associations with AD status were tested using the Wilcoxon rank-sum test. P-values were adjusted for multiple testing using the Benjamini & Hochberg (BH) method.

To identify genes associated with pseudotime, we used the median based bi-weight midcorrelation (BWC)^22^, more robust to outliers, as implemented in R-WGCNA. Genes with BH adjusted P (BHP) < 0.05 and |BWC| ≥ 0.60 were considered statistically significant. To prioritize genes showing consistent pseudotime-associated expression changes across brain datasets, we combined the region-level results with a meta-analytical approach. We retained statistically significant genes only showing concordant correlation direction across all regions. For these concordant genes, region-specific BWC coefficients were *Fisher-Z* transformed and combined using a random-effect meta-analysis as implemented in the *metafor* R package. Sampling variances were estimated as *1/(n -3)*, where *n* is the region-specific sample size. Meta-analytic effect sizes, confidence intervals, P values, and between-region heterogeneity metrics were obtained using restricted maximum likelihood estimates. Effect sizes were back-transformed to the correlation scale for reporting.

Gene Ontology (GO) enrichment analysis^23^ was performed using the R-package *clusterProfiler* (*v4.12.6*), separately for each brain region. For each region, two gene sets were tested: positively and negatively correlated genes with pseudotime (BHP < 0.05 and |BWC| ≥ 0.60). The analysis was conducted for each gene set using the three GO classes (biological process, cellular component and molecular function). The background universe comprised all genes with an Entrez identifier tested in that region. Redundant GO terms were collapsed with the *simplify* function from *clusterProfiler,* using the Want semantic-similarity measure with a cutoff of 0.7 and retaining, within each group of similar terms, the term with the smallest adjusted P value. P-values were adjusted for multiple testing using the BH method and considered statistically significantly when BHP was less than 0.05.

Enrichment analysis for cell type specific genes among the pseudotime-associated genes was performed using marker genes lists obtained as described in Piras et a al^24^, derived from the single-nucleus dataset of Mathys et al^25^ and comprising eight cell types: inhibitory neurons (In), excitatory neurons (Ex), microglia (Mic), astrocytes (Ast), pericytes (Per), oligodendrocytes (Oli), oligodendrocyte precursor cells (OPC) and endothelial cells (End).

Statistical enrichment was assessed using a hypergeometric test as implemented in the R-package *bc3net,* using as the background universe all genes tested in that specific brain region. P-values were adjusted for multiple testing using the BH method.

### Multiscale Embedded Gene co-Expression Network Analysis (MEGENA) and Key Driver Analysis (KDA)

Co-expression analysis was performed with the *MEGENA* R package^26^, using as input the same expression matrices used for pseudotime trajectories estimation. Pairwise correlations among genes were computed using Pearson’s correlation, and significant correlations after 1,000 permutations (BHP < 0.05) were retained to build a planar filtered network (PFN) using the planar maximally filtered graph (PMFG) algorithm. The PFN was subsequently decomposed into multiscale clusters (minimum module size: 50 nodes) through multiscale clustering analysis (MCA) to identify hierarchically organized co-expression modules. Module significance was evaluated by permutation testing (n = 1,000; P < 0.05), and significant hub genes (P < 0.05) were identified within each module based on intra-module connectivity (module degree). We then extracted the module eigengenes and examinated their relationship with pseudotime trajectories using linear regression, with module eigengene as the outcome and pseudotime as independent variable. For modules significantly associated with pseudotime, we performed cell type specific and GO enrichment analyses as described above, using as background universe all the genes included in coexpression modules in that specific region. All P-values were adjusted for multiple testing using the BH method.

Weighted Key Driver analysis (kDA) was conducted using the Mergeomics^27^ web tool projecting the significantly associated coexpression modules into brain specific Bayesian reference networks. Specifically, GTEx-V8 Cingulate Cortex networks were used for the ACC and PCC regions; Frontal Cortex networks for DLPFC, FP, IFG; Hippocampus for PHG; Cortex for STG and TCX; and cerebellum for CBE. Key drivers were identified using the default Mergeomics parameters: a search depth of one edge, undirected edges (*direction* = 0), equal edge weighting (*edgefactor* = 0), and a maximum subnetwork overlap of 0.33 (*maxoverlap* = 0.33). Key drivers were considered significant at FDR < 0.05.

To identify shared and brain-specific co-expression patterns (metamodules), we compared modules pairwise across all brain regions analyzed. Because *MEGENA* modules were generated independently for each brain region, module identifiers are region-specific and cannot be compared directly across regions. We therefore performed module matching based on gene membership similarity and preservation statistics, to identify modules representing analogous transcriptional programs across regions. For each brain region, we retained only modules associated with pseudotime (BHP < 0.05). Because MEGENA modules are hierarchically nested, we selected a non-redundant set by retaining, within each branch of the module hierarchy, the most significantly pseudotime-associated module; where several modules withing a branch occupied the same hierarchical level, the one with the lowest adjusted P value was retained. For every pair of brain regions, all selected modules from one region were compared with all selected modules from the other region. Pairwise module similarity was quantified using three metrics: (1) the number of shared genes; (2) the Jaccard index, defined as the size of the gene-set intersection divided by the size of the union, and (3) pairwise hypergeometric enrichment (‘*enrichment*’ function, *R-bc3net*) using as the universe background the intersection of genes detected in two regions compared, adjusting the enrichment P-values with the BH method. We additionally assessed module preservation for each module pair using the *modulePreservation* function implemented in WGCNA, with 1,000 permutations. Preservation was summarized by *Zsummary,* with *Zsummary* >10 taken to indicated strong preservation and values between 2 and 10 weak to moderate preservation. Module matches were classified as (1) ‘strong’ when they showed enrichment BHP ≤ 0.05, JI ≥ 0.20, and ≥20 overlapping genes; (2) ‘moderate’ when enrichment BHP ≤ 0.05, JI ≥ 0.10, and ≥10 overlapping genes. All remaining pairs were classified as non-matching. Module-overlap results were integrated with the independent module-preservation statistics, and downstream summaries were restricted to strong module matches showing strong preservation.

### Use of Large Language Models

Large language models (LLMs) were used to assist with language editing. All output was reviewed and verified by the authors, who take full responsibility for the content.

## Results

### Pseudotime tracks clinical and neuropathological progression and defines a conserved transcriptional signature

We inferred pseudotime trajectories for each brain region with *phenoPath*^7^ and assessed their association with clinical and neuropathological variables, for a total of 41 comparison across all brain regions. After adjusting for multiple testing across all regions and variables (BH method), 82.9% of the comparisons were significant (**Table S2**). Notably, pseudotime was significantly associated with AD diagnosis in all nine brain regions (**Fig. 1A)**, and with Braak stage was associated in seven of nine brain regions (**Fig**. **2B**).

**Figure 1.**
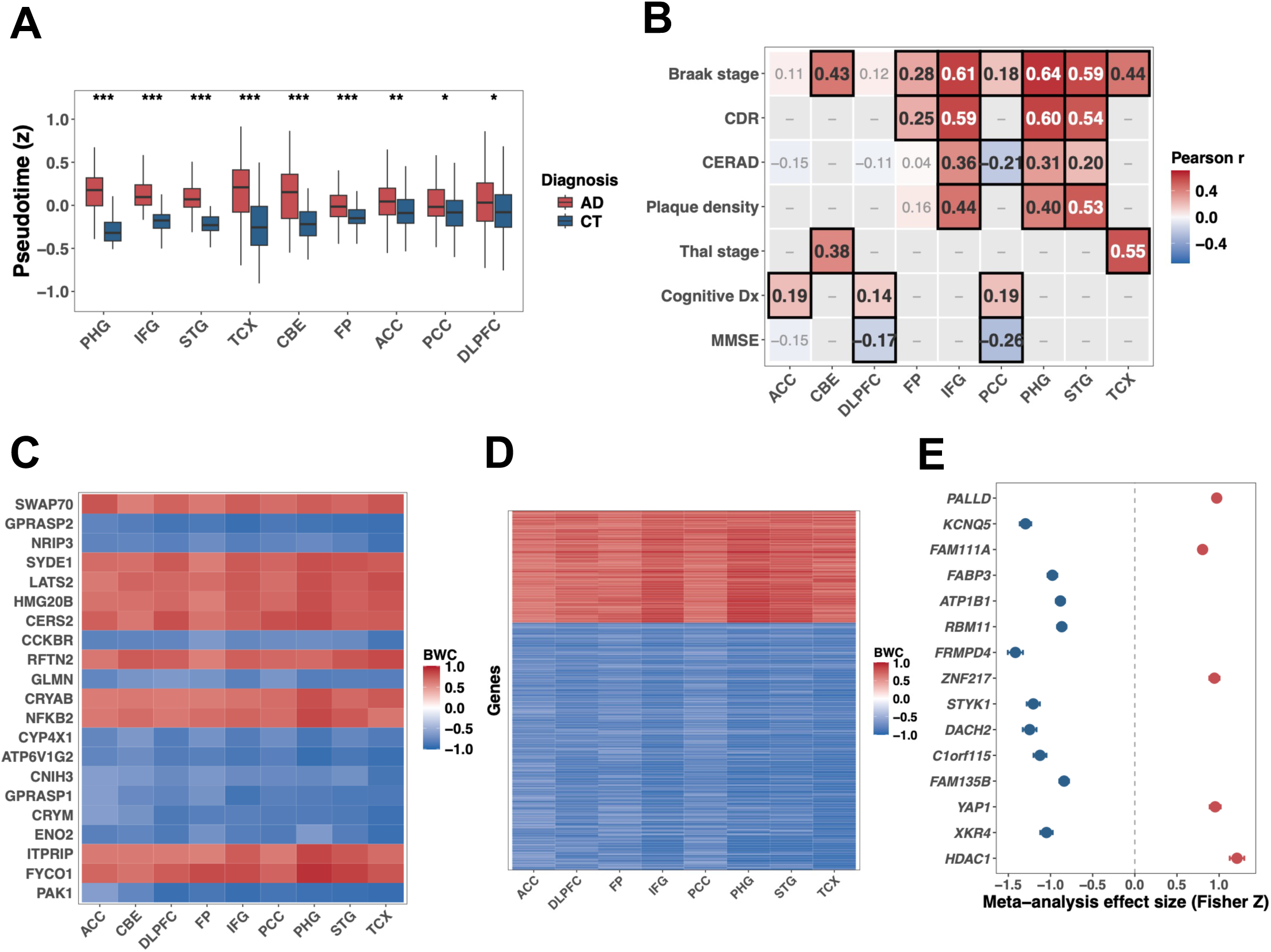
Pseudotime trajectories are associated with clinical and neuropathological progression and define a conserved transcriptional signature across nine brain regions. **(A)** Distribution of pseudotime values in AD diagnosis (AD vs control) across nine brain regions. Wilcoxon rank-sum test (*p < 0.05, **p < 0.01, ***p < 0.001). **(B)** Heatmap of Pearson correlations between pseudotime and clinical/neuropathological variables across brain regions. Significant associations (BHP < 0.05) are shown with black bordered tiles; dashes indicate missing clinical variables. Note: CERAD scoring direction differs between cohorts (ROSMAP: 1 = frequent plaques, 4 = none; MSBB: 0 = none, 3 = frequent) **(C)** Heatmap of biweight midcorrelation (BWC) with pseudotime for 21 genes with concordant associations across all nine brain regions. **(D)** Heatmap of BWC with pseudotime for 234 genes with concordant associations across eight cortical regions (excluding CBE). **(E)** Forest plot of meta-analysis effect sizes (Fisher Z-transformed) for the top 15 genes prioritized across eight cortical regions. Red and blue indicate positive and negative pseudotime associations, respectively.

**Figure 2.**
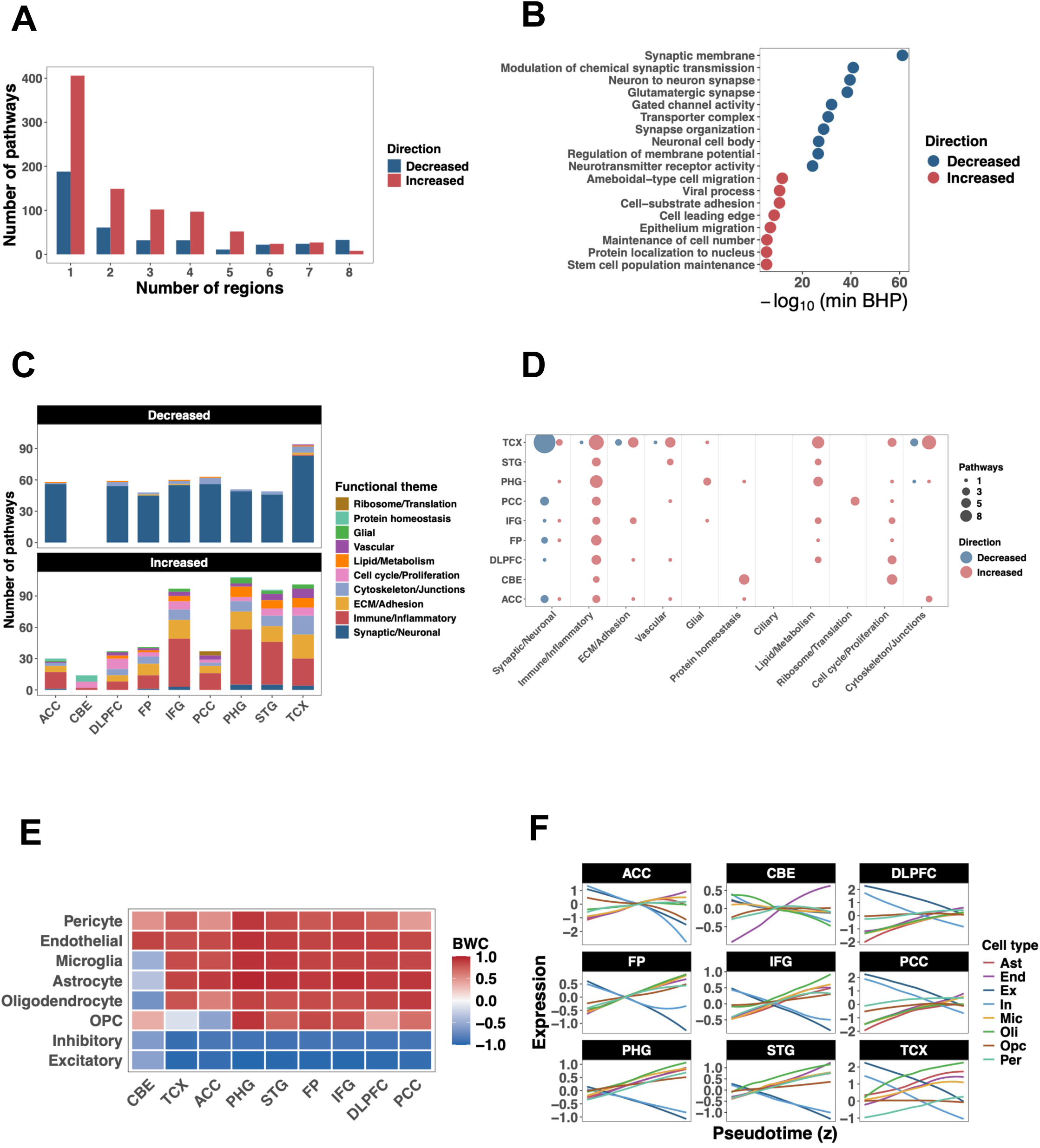
Pseudotime-associated pathways across brain regions. **(A)** Number of significantly enriched GO pathways by number of brain regions, stratified by direction (decreased or increased along pseudotime). **(B)** Top GO pathways enriched in all eight cortical regions, ranked by -log10(min BHP). Point size indicates number of regions; color indicates direction. **(C)** Stacked bar plots of pathway counts per region grouped by functional theme, shown separately for decreased (top) and increased (bottom) pathways. **(D)** Bubble plot of region-specific pathway counts grouped by functional theme. Point size indicates number of pathways; color indicates direction. **(E)** Heatmap of BWC between cell-type-specific expression signatures and pseudotime across brain regions. **(F)** Cell-type expression trajectories along pseudotime for each brain region, colored by cell type (Ast, astrocytes; End, endothelial; Ex, excitatory neurons; In, inhibitory neurons; Mic, microglia; Oli, oligodendrocytes; Opc, oligodendrocyte precursor cells; Per, pericytes).

We assessed the correlation between individual gene expression and PT trajectories, identifying between 901 (ACC) to 2,492 (PHG) significantly associated genes per region (BHP < 0.05; |BWC| ≥ 0.6) (**Table S3**). Twenty-one genes were significantly correlated with PT in a concordant direction in all nine brain regions (**Fig. 1C**) (**Tables S4**). After prioritization by meta-analysis, the top associated genes were *SWAP70*, *GPRASP2*, *NRIP3* and *SYDE1*. Negatively correlated genes included several involved in G Protein-Coupled Receptors signaling (GPCRs) (*GPRASP2*, *GPRASP1*, *CCKBR*), and vesicle-mediated transport (*PAK1*). Positively correlated genes included regulators of chromatin remodeling (*HMG20B*), protein folding (*CRYAB*), microtubule transport of autophagosomes (*FYCO1*), and TLR4-associated receptors (*RFTN2*). When CBE was excluded, which displayed a distinct transcriptional pattern relative to other regions, the number of concordant genes increased to 234, demonstrating a distinct pattern (**Table S5**; **Fig. 1D**). The top associated genes, prioritized by meta-analysis, were *PALLD*, *KCNQ5*, *FAM111A* and *FABP3* (**Figure 1E**).

### Shared synaptic and immune pathway remodeling replicates across cohorts, with the cerebellum as an exception

GO enrichment analysis of pseudotime associated genes identified 1,268 significant pathways across all brain regions. Among these, 201 were significantly enriched in five of more regions (**Table S6**), 473 in two to four regions (**Table S7**), and 594 were unique to a single region (**Table S8**). Downregulated pathways dominated the most prevalent shared set (80% of pathways enriched in all eight cortical regions), whereas upregulated pathways were increasingly prevalent among less broadly shared associations (83% at five regions, 71% at two regions) (**Fig. 2A**). Among the 41 pathways detected in all eight cortical regions, 33 were downregulated and 8 upregulated (top pathways showed in **Fig. 2B**). The downregulated set was almost exclusively synaptic and neuronal, led by synaptic membrane (min BHP = 6.7×10^-62^), modulation of chemical synaptic transmission (1.2 × 10^-41^), glutamatergic synapse (3.4 × 10^-39^), regulation of membrane potential (3.2 × 10^-27^), neurotransmitter receptor activity (6.5×10^-25^), and GABAergic synapse (2.0 × 10^-21^). The eight top upregulated pathways included ameboidal-type cell migration (1.9 × 10^-12^), viral process (2.4 × 10-11), cell-substrate adhesion (2.6 × 10^-11^), and cell leading edge (4.1 × 10^-09^) (**Fig. 2B**).

We grouped the pathways by functional theme (keyterms used for classification are reported in **Table S9**), finding that synaptic/neuronal processes represented the largest category and were decreased along pseudotime, while immune/inflammatory pathways were predominantly increased (**Fig. 2C, Table S10**). This pattern was consistent across regions, though the relative contribution of each theme varied: TCX and IFG showed the strongest synaptic decline, PHG and FP the most ECM/adhesion enrichment, and PCC unique upregulation of ribosomal and translational pathways (**Fig. 2D**). CBE showed a different pattern, with only 18 region-specific pathways detected, which 17 were upregulated, enriched for protein refolding, chaperone activity, and chromatin structure rather than the neuronal decline and immune activation characterizing cortical regions (**Fig. 2C** and **Fig. 2D**).

### Cell-type deconvolution links progression to neuronal loss and astrocyte-led glial activation

We assessed cell-type specific expression changes along pseudotime for eight major brain cell types (**Fig. 2E** and **Fig. 2F**; **Table S11**). Consistent with the neurodegeneration process in AD, excitatory and inhibitory neuron signatures showed a strong decrease along pseudotime across all nine brain regions (BWC range: -0.57 to -0.99 for excitatory neurons, -0.62 to -0.94 for inhibitory neurons). In cortical regions, Ast, End, Mic, Oli, OPCs, and Per increased across pseudotime, with Ast and Mic showing the strongest positive association (BWC 0.96 and 0.92, respectively). OPC signatures were positive in most cortical regions but decreased in ACC (BWC = -0.56) and were non-significant in TCX. CBE shows a different pattern with negative association for Oli, Mic, Ast, while Per, End, Ex and In followed the trend observed in cortical regions.

### Multiscale Embedded Gene co-expression Network Analysis (MEGENA)

We performed multiscale gene-coexpression network analysis using MEGENA^26^, identifying 874 modules significantly associated with pseudotime across the 9 brain regions (**Table S12**). The number of significant modules ranged from 78 (CBE) to 113 (DLPFC). The results highlighted differential involvement of neuronal, immune, glial, and extracellular matrix-related programs across regions. To identify conserved cross-regional transcriptional programs associated with the coexpression networks, we grouped the modules into metamodules using a combination of gene overlap, Jaccard index (JI), hypergeometric enrichment, and module preservation statistics. We detected 83 unique strong correspondences (enrichment BHP < 0.05, JI ≥ 20, overlap ≥ 20, and strong module preservation) (**Table S13**), grouped into six metamodules (MM1 – MM6) (**Table S14** and **Table S15**). The 43 modules assigned to metamodules showed a predomincance of positive pseudotime associations: 69.8% were positively associated, whereas 30.2% were negatively associated (**Fig. 3A**). Positively associated modules were mostly enriched for glial cell types, whereas negatively associated modules were exclusively enriched for excitatory neurons. Excitatory neuronal modules represented the largest proportion of preserved matches (30.2%), followed by microglial (25.6%) and oligodendrocyte modules (25.6%). Additionally, excitatory and oligodendrocyte modules exhibited higher preservation statistics than microglial modules (mean *Zsummary* = 35.2 and 35.0 versus 27.2, respectively), whereas microglial modules displayed the highest fold-enrichment of gene overlap between regions (mean fold-enrichment = 4.57 vs 2.00 for excitatory neurons).

**Figure 3.**
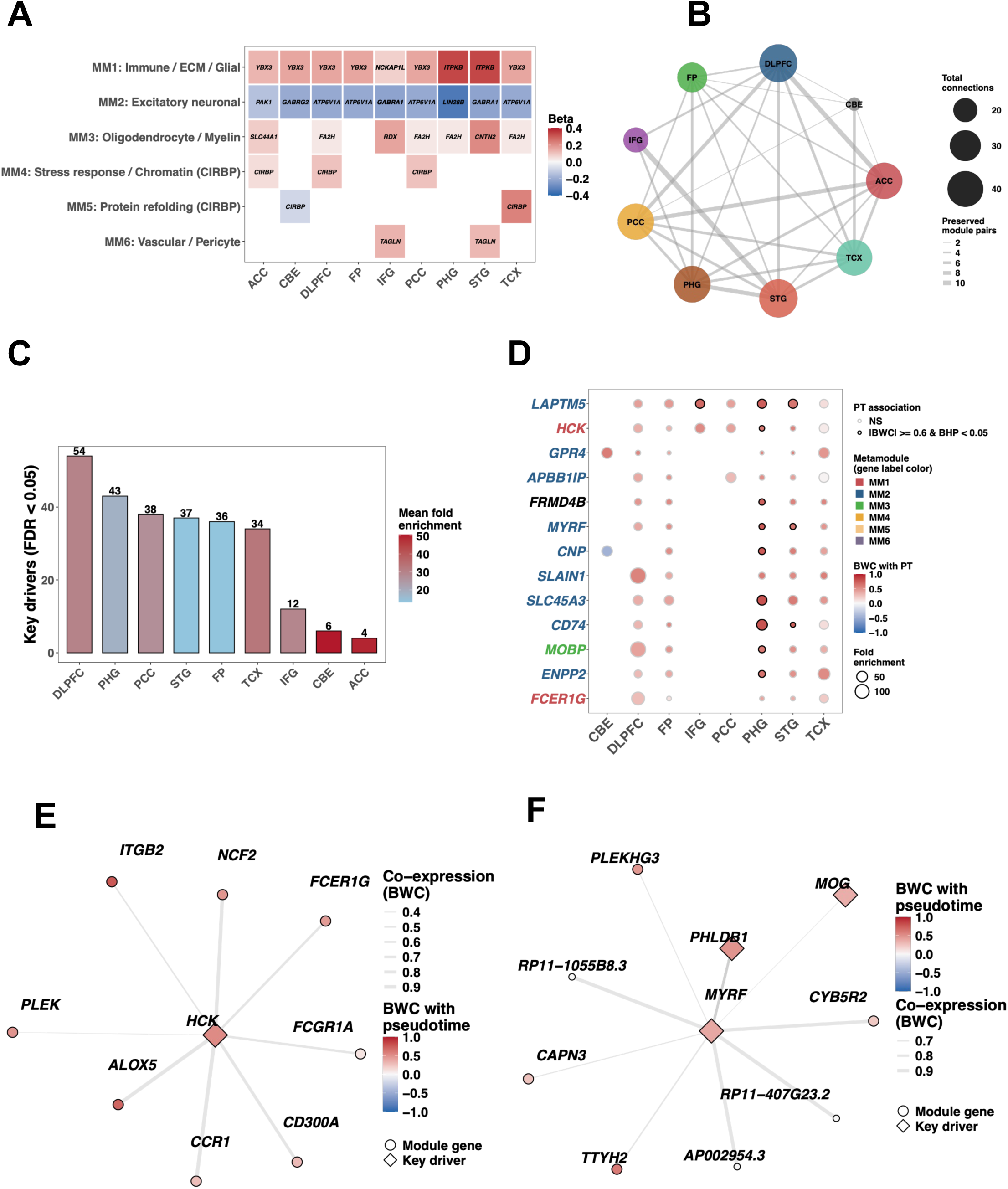
Conserved metamodules and cross-regional key drivers of AD progression detected by pseudotime. **(A)** Tile plot of six metamodules (MM1-MM6) across brain regions. Fill color indicates pseudotime association strength (beta coefficient); labels indicate hub genes for each region-module pair. **(B)** Network plot of regional connectivity based on preserved module correspondences. Node size reflects total connections; edge width reflects number of preserved module pairs between regions. **(C)** Number of significant key drivers (FDR < 0.05) per brain region. Bar color reflects mean fold-enrichment. **(D)** Key driver genes detected in five or more brain regions. Point size indicates fold-enrichment; fill color indicates BWC with pseudotime; point border distinguishes significant (|BWC| ≥ 0.6 and BHP < 0.05) from non-significant pseudotime associations. Gene label color indicates metamodule assignment (MM1, red; MM2, blue; MM3, green; MM4, amber; MM5, gold; MM6, purple). **(E)** Co-expression subnetwork of *HCK* in IFG. Node fill indicates BWC with pseudotime; node shape distinguishes key drivers (diamond) from module genes (circle); edge width indicates co-expression strength (BWC). **(F)** Co-expression subnetwork of MYRF in DLPFC, with the same encoding as panel E.

Two metamodules were detected across all nine brain regions. MM1 comprised 15 modules and was characterized by microglial and astrocyte-enriched modules, all positively associated with pseudotime (mean β = 0.155). GO analysis showed enrichment for cytokine production, immune receptor activity, extracellular matrix organization, cell adhesion, and wound healing, with hub genes including *NCKAP1L*, *ITPKB*, and *YBX3*, and key drivers *GPR4*, *HCK*, and *ANXA2*. MM2 comprised 12 excitatory neuronal modules and was negatively associated with pseudotime (mean β = −0.215). These modules were enriched for modulation of chemical synaptic transmission, regulation of membrane potential, and synaptic membrane organization, with hub genes *GABRA1*, *GABRG2*, *LIN28B*, and *PAK1*. *LIN28B* was the hub gene of the module showing the strongest negative association with pseudotime in FP (β = −0.342). MM3 included 9 oligodendrocyte-enriched modules across seven cortical regions (excluding FP and CBE) and was positively associated with pseudotime (mean β = 0.077). These modules were enriched for oligodendrocyte differentiation, ensheathment of neurons, and myelin sheath, with hub genes *MOG*, *FA2H*, *CNTN2*, and *RDX*. The remaining metamodules were associated with more restricted transcriptional programs. MM4 included 3 modules from ACC, DLPFC, and PCC (hub gene: CIRBP), with enrichment for nucleosome assembly and response to misfolded proteins. MM5 included 2 modules from CBE and TCX, also with CIRBP as hub gene, with enrichment for protein refolding and chaperone activity. MM6 included 2 pericyte-enriched modules from IFG and STG centered on (hub gene: TAGLN), enriched for extracellular matrix organization.

Analysis of regional connectivity identified STG, DLPFC, and PHG as the most highly connected regions within the preserved module network, with 24, 23, and 23 connections, respectively (**Fig. 3B**). CBE showed the lowest connectivity (7 connections), with most preserved relationships involving TCX. The most frequently connected regional pairs were IFG– STG (11 module pairs) and ACC–DLPFC (10 pairs), followed by ACC–PCC and PHG–STG (9 pairs each).

### Cross-regional key driver analysis highlights immune and oligodendrocyte regulators

Using undirected wKDA, we identified 264 significant key drivers (FDR < 0.05) across 35 modules, corresponding to 76 unique genes (**Table S16**). The number of key drivers varied across regions, with DLPFC (54), PHG (43), and STG (37) showing the highest counts, while CBE (6) and ACC (4) had the fewest (**Fig. 3C**). Two genes were significant key drivers in seven of nine regions: *HCK* (hematopoietic cell kinase; mean fold-enrichment = 20.5) and *LAPTM5* (lysosomal protein transmembrane 5; mean fold-enrichment = 19.7), both predominantly assigned to MM1 (immune/glial) modules and positively correlated with pseudotime (mean BWC = 0.42 and 0.53, respectively) (**Fig. 3D**). Additional 11 genes were key drivers in five or more regions, associated with two functional axes. The first included immune/microglial genes, such as GPR4 (6 regions), APBB1IP (6 regions), CD74 (5 regions), and FCER1G (5 regions), predominantly mapping to MM1 modules. The second comprised oligodendrocyte/myelin-associated genes, such as MOBP (mean fold = 38.3), SLC45A3 (24.8), ENPP2 (25.6), MYRF (14.0), and CNP (22.0), each significant in five regions and predominantly assigned to MM3 modules. SLAIN1, associated with microtubule dynamics, showed the highest mean fold-enrichment among broadly replicated drivers (46.2, five regions). Finally, FRMD4B was a broadly replicated key driver (5 regions) with a strong positive pseudotime correlation (mean BWC = 0.57). Networks for HCK and MYRF are reported in **Figs 3E** and **3F**.

## Discussion

We leveraged pseudotime on harmonized bulk RNA sequencing to capture a reproducible axis of AD progression across nine brain regions. Mukherjee et al. previously demonstrated the feasibility of reconstructing AD-related pseudotime from AMP-AD bulk transcriptomic data from dorsolateral prefrontal and temporal cortex, reporting associations with Braak, CERAD score, and cognitive diagnosis^14^. Our findings confirm those observations using *phenoPath*, while extending the analysis to seven additional regions and integrating 2,106 samples from three AMP-AD cohorts across nine brain regions.

The inferred trajectories were associated with diagnosis and with clinical or neuropathological severity, remaining significant in 34 of 41 comparisons after multiple-testing correction, including AD diagnosis in all nine regions and Braak stage in seven of nine. Pseudotime should therefore be viewed as a molecular correlate of disease burden, capturing robust features of AD progression rather than replacing neuropathological staging.

Cross-region pathway analysis revealed a conserved cortical progression characterized by declining neuronal and synaptic programs and increasing inflammatory and extracellular-matrix-related processes. Among pathways enriched in all eight cortical regions, 80% were downregulated, dominated by synaptic and neuronal processes including synaptic membrane, glutamatergic signaling, and neurotransmitter receptor activity. Upregulated pathways became increasingly prevalent among less broadly shared associations, suggesting that while synaptic decline is the most conserved transcriptional feature of AD progression, immune and tissue remodeling programs show greater regional specificity. These findings are consistent with previous transcriptomic meta-analyses^20,28^ and single-cell studies of AD^25,29–38^ demonstrating a coordinated transition from neuronal homeostasis toward inflammatory and stress-associated states during progression. Importantly, these coordinated changes were shared across cortical regions rather than representing isolated regional signatures.

The cerebellum represented the main exception, showing fewer disease-associated pathways, weaker neuronal decline, and enrichment for protein folding and chaperone activity. Only 18 region-specific pathways were detected in CBE (17 upregulated), in contrast to the neuronal decline and immune activation characterizing cortical regions. This observation is consistent with neuropathological studies describing the relative resistance of the cerebellum to classical AD pathology compared with vulnerable cortical regions^39^. However, our findings extend previous observations by suggesting that cerebellar resistance is associated not only with reduced pathological burden but also with a distinct transcriptional response during aging and AD progression. The enrichment of protein-folding and chaperone-related pathways may reflect region-specific mechanisms of stress adaptation or cellular resilience, although functional studies will be required to determine whether these pathways contribute to cerebellar protection. Cell-type enrichment analysis provided complementary evidence for this progression, showing a coordinated decline of neuronal signatures and increase of glial-, vascular-, and oligodendrocyte-associated signatures along pseudotime. Excitatory and inhibitory neuron signatures showed strong negative correlation with pseudotime across all nine regions, while astrocytes and microglia showed the strongest positive associations in cortical regions. CBE followed a distinct pattern, with microglia, astrocytes, and oligodendrocytes showing negative pseudotime associations, while only endothelial cells and pericytes maintained positive associations as observed in the cortical regions. These trends are consistent with cell-resolved AD studies^25,29–31^ and highlight the contribution of diverse cellular programs, including myelin-associated processes, to AD-related transcriptional changes^4,40–53^. In particular, the progressive increase in oligodendrocyte-associated signatures supports emerging evidence that myelin and white-matter-related processes represent active components of AD pathophysiology rather than merely secondary consequences of neuronal degeneration^4,40–53^. Previous transcriptomic and cell-resolved studies have highlighted alterations in oligodendrocyte states, myelin-associated genes, and lipid metabolism during AD progression, suggesting that glial responses contribute to disease evolution alongside neuronal dysfunction. Our findings extend these observations by showing that these cellular programs are coordinated along a molecular trajectory shared across multiple brain regions.

Our cross-region metamodule analysis indicates that AD progression is characterized by a conserved transcriptional architecture shared across anatomically distinct brain regions. We identified six metamodules from 83 unique strong module correspondences, with two dominant program, immune/glial (MM1, 15 modules) and excitatory neuronal (MM2, 12 modules), spanning all nine brain regions. A third conserved program, oligodendrocyte/myelin-associated (MM3, 9 modules), was present in seven cortical regions. Excitatory neuronal and oligodendrocyte modules exhibited higher preservation statistics than microglia modules, while microglial modules showed the highest fold-enrichment of gene overlap between regions, suggesting that immune programs, though less preserved in terms of co-expression structure, are more tightly coordinated in gene composition across regions. Previous transcriptomic studies have consistently identified immune activation, synaptic dysfunction, and cellular remodeling as major molecular features of AD; however, these processes have often been investigated independently within single regions. By integrating region-specific co-expression networks, we show that these biological programs are organized into highly preserved regulatory structures, suggesting that AD progression reflects coordinated alterations of shared molecular pathways rather than independent regional transcriptional changes. The conserved increase of glial-associated metamodules is consistent with extensive evidence linking microglial activation, inflammatory signaling, and extracellular matrix remodeling to AD progression, while the conserved decline of neuronal-associated metamodules agrees with previous reports of progressive synaptic and neuronal dysfunction. Interestingly, oligodendrocyte-associated programs were also preserved across regions and increased along pseudotime, suggesting that myelin-related changes may represent active remodeling or compensatory responses rather than solely progressive loss, consistent with emerging evidence of dynamic glial responses during neurodegeneration. Together, these findings support a model in which AD progression involves conserved cellular programs, including neuronal dysfunction, immune activation, and myelin-associated remodeling, embedded within region-specific transcriptional contexts.

Key-driver analysis identified candidate regulators associated with conserved pseudotime-associated network architecture across brain regions. Among the 76 unique key drover genes identified across 35 modules, *HCK* and *LAPTM5* emerged as the most broadly replicated, each detected in seven of nine brain regions and associated with immune-related transcriptional programs (MM1). *LAPTM5* shows established expression in microglial populations and it is involved in lysosomal and immune signaling pathways; therefore, the progressive increase of *LAPTM5* along pseudotime supports the activation of conserved microglial programs during AD progression. *HCK*, a Src-family kinase expressed in microglia and myeloid cells, showed a similar positive pseudotime association, suggesting a conserved regulatory role in neuroimmune activation. Additionally, broadly replicated immune key drivers included *GPR4* (6 regions), a pH-sensing GPCR with emerging associations to neuroinflammation and one of the few key drivers detected in the cerebellum, and *CD74* (5 regions), a component of MHC class II antigen presentation. The recurrent identification of these immune regulators across independent brain regions highlights their potential relevance as regulatory nodes linking immune signaling and AD-associated transcriptional remodeling. In parallel, several key drivers associated with oligodendrocyte and myelin-related modules, including *MYRF*, *CNP*, *MOBP*, and *SLC45A3*, were identified across five regions each. MYRF, the master transcription factor for oligodendrocyte myelination, showed the strongest pseudotime correlation among myelin-associated key drivers, while *MOBP* showed the highest fold-enrichment. The positive association of these genes with pseudotime in cortical regions suggests that myelin-associated programs are not simply lost during AD progression but may undergo active remodeling, potentially reflecting compensatory or repair-associated responses. Notably, *CNP* displayed an opposite trajectory in the cerebellum, decreasing along pseudotime and further supporting the distinct molecular evolution of this region compared with vulnerable cortical areas. Together, the convergence of immune-associated (*LAPTM5*, *HCK*, *GPR4*, *CD74*, *FCER1G*) and oligodendrocyte-associated (*MYRF*, *CNP*, *MOBP*, *SLC45A3*) key drivers suggest that coordinated neuroimmune activation and myelin remodeling represent conserved regulatory features of AD progression.

Although these findings do not establish direct causal relationships, the recurrent identification of key drivers across anatomically distinct regions highlights conserved regulatory nodes underlying AD-associated transcriptional changes. These results extend previous observations by suggesting that AD progression is characterized not only by neuronal dysfunction and inflammatory activation but also by coordinated remodeling of glial and myelin-associated regulatory programs.

Several limitations should be considered when interpreting these findings. Bulk RNA sequencing averages expression across heterogeneous cell populations, and although deconvolution improves biological interpretation, it cannot fully resolve changes in cellular composition, cell-type-specific regulation, or cellular states. In addition, pseudotime reconstructs a latent molecular trajectory from cross-sectional post-mortem samples and should not be interpreted as a direct longitudinal progression. wKDA was performed using undirected edges in the reference Bayesian Network; future analyses using directed edges may refine the distinction between upstream regulators and downstream targets. Finally, the absence of APOE- and sex-stratified analyses limits the ability to assess genetically or disease-specific regulatory differences. Furthermore, the lack of validation across alternative trajectory inference methods and independent cohorts limits the assessment of the robustness and generalizability of our findings.

Future studies should evaluate trajectory stability across alternative inference methods and independent cohorts, while integrating single-cell, single-nucleus, and spatial transcriptomic datasets to determine whether conserved metamodules correspond to specific cellular states and anatomical contexts. Functional studies of recurrent candidates, including *LAPTM5*, *HCK*, *GPR4*, and oligodendrocyte-associated regulators such as *MYRF* and *MOBP*, will be required to determine their contribution to conserved AD-associated transcriptional programs.

Rather than identifying isolated transcriptional signatures, this study demonstrates that molecular features associated with AD progression converge on conserved co-expression programs across brain regions while retaining region-specific characteristics. The convergence of pathway analysis, cell-type-associated signatures, network preservation, and key-driver inference highlights recurrent regulatory nodes involved in neuroimmune activation and myelin-associated remodeling, including *LAPTM5*, *HCK*, *GPR4*, and oligodendrocyte-related drivers such as *MYRF*, *CNP*, and *MOBP*. These findings support a model in which AD progression is driven by conserved regulatory programs that integrate neuronal dysfunction, immune activation, and glial remodeling across anatomically distinct brain regions.

## Acknowledgments

The results published here are in whole or in part based on data obtained from The AD Knowledge Portal (https://doi.org/10.7303/9618241). Data generation was supported by the following NIH grants: P30AG10161, P30AG72975, R01AG15819, R01AG17917, R01AG036836, U01AG46152, U01AG61356, U01AG046139, P50 AG016574, R01 AG032990, U01AG046139, R01AG018023, U01AG006576, U01AG006786, R01AG025711, R01AG017216, R01AG003949, R01NS080820, U24NS072026, P30AG19610, U01AG046170, RF1AG057440, and U24AG061340, and the Cure PSP, Mayo and Michael J Fox foundations, Arizona Department of Health Services and the Arizona Biomedical Research Commission. We thank the participants of the Religious Order Study and Memory and Aging projects for the generous donation, the Sun Health Research Institute Brain and Body Donation Program, the Mayo Clinic Brain Bank, and the Mount Sinai/JJ Peters VA Medical Center NIH Brain and Tissue Repository. Data and analysis contributing investigators include Nilüfer Ertekin-Taner, Steven Younkin (Mayo Clinic, Jacksonville, FL), Todd Golde (University of Florida), Nathan Price (Institute for Systems Biology), David Bennett, Christopher Gaiteri (Rush University), Philip De Jager (Columbia University), Bin Zhang, Eric Schadt, Michelle Ehrlich, Vahram Haroutunian, Sam Gandy (Icahn School of Medicine at Mount Sinai), Koichi Iijima (National Center for Geriatrics and Gerontology, Japan), Scott Noggle (New York Stem Cell Foundation), Lara Mangravite (Sage Bionetworks).

## Author contributions

FE, SS, MN, and ISP performed data analysis. FE, MJH, and ISP wrote the manuscript. MHJ and ISP conceptualized the study. All authors read and approved the final manuscript.

